# Repetitive mild traumatic brain injury impairs resting state fMRI connectivity and alters protein profile signaling networks

**DOI:** 10.1101/2022.09.21.508917

**Authors:** Sakthivel Ravi, Marangelie Criado-Marrero, Daylin Barroso, Isadora M Braga, Mackenzie Bolen, Uriel Rubinovich, Gabriela P. Hery, Matteo M Grudny, John Koren, Stefan Prokop, Marcelo Febo, Jose Francisco Abisambra

**Affiliations:** Center for Translational Research in Neurodegenerative Disease, University of Florida, Gainesville, FL 32610, USA; Department of Neuroscience, University of Florida, Gainesville, FL 32610, USA; McKnight Brain Institute, University of Florida, Gainesville, FL 32610, USA; Department of Psychiatry, University of Florida, Gainesville, FL 32610, USA; Department of Pathology, University of Florida, Gainesville, FL 32610, USA; Fixel Institute for Neurological Diseases, University of Florida, Gainesville, FL 32610, USA; Brain Injury Rehabilitation and Neuroresilience (BRAIN) Center, University of Florida, Gainesville, FL 32610, USA

**Keywords:** CHIMERA, repetitive mild TBI, diffusion tensor imaging, microglia, resting state fMRI, optic tract, thalamus

## Abstract

Repetitive mild traumatic brain injury (rmTBI) is a leading and severe threat to cognition that often goes undiagnosed. A major challenge in developing diagnostics and treatments for the consequences of rmTBI is the fundamental knowledge gaps that explain how rmTBI promotes brain dysfunction. It is both critical and urgent to understand the neuropathological and functional consequences of rmTBI to develop effective therapeutic strategies. In this study, we sought to define the extent of altered brain functional connectivity (FC) and expression of neuropathological markers after rmTBI. We performed two rmTBI (2x 0.6□J impacts 24□h apart) in male and female C57BL/6J wild-type (WT) (~2.5-3mo) mice using closed head injury model of engineered rotational acceleration (CHIMERA) or sham procedures. At 5-6 days post-injury (dpi), we measured changes in brain volume and FC using T2-weighted images, resting-state functional MRI (rsfMRI), and graph theory analyses. We used diffusion tensor imaging (DTI) to assess microstructural changes in white matter tracts. In addition, at 7dpi, we measured changes in Iba1 and GFAP to determine the extent of gliosis. The expression of disease-associated protein markers in grey and white matter regions were evaluated using the NanoString-GeoMx digital spatial protein profiling (DSP) platform. The rsfMRI data revealed aberrant changes in connectivity such as node clustering coefficient, global and local efficiency, participation coefficient, eigenvector centrality, and betweenness centrality in thalamus and other key brain regions that process visual, auditory, and somatosensory information. In addition, DTI revealed significantly decreased fractional anisotropy (FA) and axial diffusivity in the optic tract. Also, mean, radial, and axial diffusivity (L1) were significantly increased in the hippocampus. DSP revealed that phospho-serine 199 tau (pS199) as well as glial markers such as GFAP, cathepsin-D, and Iba1 were significantly increased in the optic tract. In thalamic nuclei, the neuroinflammatory marker GPNMB was increased significantly, and the cell proliferation marker Ki-67 was decreased in the rmTBI group. Our data suggest that rmTBI significantly alters brain functional connectivity and causes a profound inflammatory response in gray matter regions, beyond chronic white matter damage.

## 1. Introduction

Repetitive mild traumatic brain injuries (rmTBI) are the most frequently diagnosed form of head injury in the United States [1–4]. These repetitive brain injuries result from multiple and subsequent impacting blows to the head, and they are common in subjects engaging in contact sports as well as by active military personnel. Moreover, evidence suggests that TBIs, and in particular rmTBIs, significantly increase risk for numerous neurodegenerative disorders, including chronic traumatic encephalopathy (CTE) and Alzheimer’s disease (AD), among other AD-related disorders (ADRD) [5–9]

The mechanisms linking rmTBIs and the pathogenesis of neurodegenerative disorders remain unknown. However, advances in monitoring both the anatomical and molecular changes that occur after rmTBI suggest links between TBI-induced brain alterations are detectable. For example, diffuse tensor imaging (DTI) in MRI identifies shears in white matter tracts that result in diffuse axonal injury (DAI), which is a typical sign of rmTBI [10]. In addition, increased expression of the microglial marker, Iba1 (ionized calcium-binding adapter molecule 1) and astrocytic marker, GFAP (glial fibrillary acidic protein) reflect gliosis response after injury. Though these network and molecular changes reveal considerable mechanistic information and can be found in TBIs of different severity, distinct markers of rmTBI are not clearly established.

In this study, we used functional brain imaging and digital spatial protein profiling to identify markers of altered neural function following rmTBI. Given that rmTBI causes rotational damage that ruptures white matter axonal tracts, we examined the effects of rmTBI on regions that possess high and low amount of white matter (optic tract and commissures and thalamic nuclei, respectively). Additionally, due to the purported link between TBI and cognitive damage, we examined brain structures highly involved in ADRDs (hippocampus and cortex). We identified region-specific abnormalities resulting from rmTBI. These distinct imaging and protein surrogates of brain damage strongly suggest that, though the global consequences of rmTBIs are unpredictable, the local region-specific effects are consistent and are reflected at the gross structural and fine molecular levels. This information reveals novel imaging and protein biomarkers that could help determine outcomes and inform mechanistic knowledge gaps.

## 2. Materials and Methods

### 2.1 Animals

Male and female C57BL/6J (WT) mice were purchased from the Jackson Laboratory (Bar Harbor, ME, USA). Two to five mice of same-sex groups were housed in standard room conditions with 12 h light and dark cycle and allowed to access free food and water ad libitum. All the animal procedures were approved by the Institutional Animal Care and Use Committee (IACUC), University of Florida.

### 2.2 Repetitive mild traumatic brain injury (rmTBI) by CHIMERA

The rmTBI procedure was followed as suggested by Namjoshi et al. 2014[11]. 2.5-3 months old male and female WT mice were subjected to two mild closed head injuries (0.6 J) at 24h intervals using the Closed-Head Impact Model of Engineered Rotational Acceleration (CHIMERA) impactor. Briefly, mice were anesthetized with isoflurane (induction 3-4% and maintenance 1-2.5%), and lubricating eye ointment was applied to prevent corneal drying. Meloxicam (10 mg/kg) was administered subcutaneously right before injury for two days to alleviate pain. Animals were positioned supine in the holding bed of the impactor. The head was aligned in a flat position over a hole in the head plate to receive piston strikes closer to the bregma. After impact, animals were transferred immediately to the recovery chamber pre-warmed at 38°C and monitored until fully ambulatory. Sham mice were exposed to all these procedures, except for the impact. At 7dpi, transcardial perfusions were performed with sterile ice-cold 0.9 % saline and the brain tissues were harvested. One half of the hemibrain was micro-dissected and stored at −80°C ultra-deep freezer for further analysis. The other half of the hemibrain was fixed in 10% neutral buffered formalin for further histological analysis.

### 2.3 T2 and resting-state functional Magnetic Resonance Imaging (rsfMRI) data acquisition

Anatomical T2-weighted, diffusion weighted and rsfMRI scans were collected between 5-6 dpi. Sham and rmTBI mice were anesthetized with a mixture of isoflurane (Induction 4%, maintenance: 1%) and medical grade air (0.1-0.15L/min) and positioned prone on a custom-built holder with bed, ports for warm water flow, a bite bar and a nose port for gas anesthetic delivery. A respiratory pad was placed underneath the abdomen to monitor respiration throughout the entire imaging session. The core body temperature of the mice was maintained at 36-37°C by placing the circulating warm water setup connected to the water bath. Lubricating eye ointment was applied to prevent corneal drying, and a radiofrequency (RF) coil was placed on top of the head. The images were acquired using an 11.1T MRI scanner (Magnex Scientific Ltd, Oxford, UK) with high power gradient sets (RRI BFG-240/120-S7; maximum gradient strength of 1000 mT/m at 325 Amps and a 200 ms rise time; RRI, Billerica, MA). The MRI system was controlled by a Bruker Paravision 6.01 console (Bruker BioSpin, Billerica, MA). A custom-made 2.0 cm x 2.5 cm quadrature RF surface transmit/receive coil tuned to 470.7 MHz (1H resonance) was used for B1 excitation and signal detection (RF engineering lab, Advanced Magnetic Resonance Imaging and Spectroscopy Facility, Gainesville, FL). A T2-weighted Turbo Rapid Acquisition with Refocused Echoes (TurboRARE) sequence was acquired with the following parameters: effective echo time (TE) = 41.42 ms, repetition time (TR) = 4s, RARE factor =16, number of averages = 12, the field of view of 15 mm x 15 mm and 0.9 mm thick slice, a data matrix of 256 × 256 and 14 interleaved ascending coronal (axial) slices covering the entire brain from the rostral-most extent of the anterior frontal cortical surface, caudally towards the upper brainstem and cerebellum. The resting-state functional images were collected using a single-shot spin-echo echo planar imaging sequence with the following parameters: TE = 15 ms, TR = 2 s, 600 repetitions, the field of view = 15 × 15 mm and 0.9 mm thick slice, and a data matrix of 64 × 48 with 14 interleaved ascending coronal slices in the same space as the T2 anatomical. Two additional single repetition SE-EPI scans were collected, one with phase encode gradient lobes (‘PE blips’) collected along the positive gradient direction and the other collected with PE blips reversed to the negative PE direction (Bruker Paravision binary method for distortion correction is kindly provided by Dr. Matthew Budde, Medical College of Wisconsin).

### 2.4 rsfMRI image pre-processing

Functional connectivity of rmTBI mice brain was analyzed by graph theoretical analysis, an approach highly used to assess novel biomarkers of brain function in neurological diseases and aging [12]. SE-EPI distortions were corrected using TOPUP in FSL [13]. The pair of opposite phase-encode blip scans were used to estimate the susceptibility-induced off-resonance field, which is used to unwarp subsequent EPI volumes [13]. Image pre-processing and analysis was carried via in-house UNIX terminal integrating the following tools: Analysis of Functional NeuroImages (AFNI)[14], FMRIB Software Library (FSL)[15–17] and Advanced Normalization Tools (ANTs) [18,19]. Anatomical and distortion-corrected functional scan masks outlining mouse brain boundaries were generated in MATLAB using Three-Dimensional Pulsed Coupled Neural Networks (PCNN3D) [20] and then manually edited. Then, 3dDespike in AFNI was used to remove time series spikes and 3dvolreg for image volume alignment. Preprocessed scans were cropped and a high-pass temporal filter (<0.009Hz) was used (3dTproject) to remove slow variations (temporal drift) in the fMRI signal. Independent component analysis (ICA) decomposition was then applied using Multivariate Exploratory Optimized Decomposition into Independent Components (FSL MELODIC version 3.0) to preprocessed scans to assess noise components in each subjects’ native space prior to spatial smoothing and registration. The resulting components were assessed, and in most cases all components contained noise-related signal along brain edges, in ventricular voxels, and large vessel regions. These components were suppressed using a soft (‘non-aggressive’) regression approach, as implemented in FMRIB Software Library (FSL 6.0.3) using fsl_regfilt [15]. A low-pass filter (>0.12Hz) and spatial smoothing (0.4mm FWHM) was then applied to the fMRI scans.

Preprocessed anatomical and fMRI scans were aligned to a parcellated mouse common coordinate framework (version 3, or CCFv3) template [21]. Bilateral region of interest (ROI)-based nodes (64 total) was created with the guidance of the annotated CCFv3 parcellation and using tools in ITKSNAP and FSL. In ITKSNAP, the template along with the overlaid parcellation were used to find the left hemisphere voxel coordinates for each of the nodes included in this study, which were distributed as evenly as possibly without overlap across the mouse brain template. The node coordinates were positioned in subregions of all areas for example the optic tract and thalamus. We created 0.6 mm diameter spheric nodes in the template space (resolution: 0.05mm) centered on the voxel coordinates. The right hemispheric representations of the same nodes were then created to complete left and right representations for each node. For subject-to-atlas registration, fMRI scans are upsampled from a native space 0.234×0.3125 x0.9 mm resolution (spatially smoothed at 0.4 mm FWHM) to a downsampled template resolution of 0.1mm^3^. Thus, the spheric nodes, in effect, were designed to sample fMRI signals at a native space slightly over single-voxel resolution.

Anatomical scans were cropped and N4 bias field correction [22] applied to T2 images to correct intensity variations due to RF field inhomogeneities [23]. The extracted brain maps were linearly registered to the mouse template using FSL linear registration tool (FLIRT)[15], using a correlation ratio search cost, full 180-degree search terms, 12 degrees of freedom and trilinear interpolation. The linear registration output was then nonlinearly warped to the template space using ANTs (antsIntroduction.sh script). Anatomical-to-atlas linear and nonlinear transformation matrices were applied to fMRI scans at a later stage. Brain extraction using a mask (see above) was first applied to fMRI scans and the cropped scans were then aligned to their respective higher resolution anatomical scans. Timeseries functional images were split into 600 individual volumes and the first in the series was linearly aligned to the anatomical scan using FLIRT (same parameters as above, except 6 degrees of freedom was used in this step). ANTs (antsRegistrationSyNQuick.sh script) was used to warp the lower resolution functional images to their structural (using a single stage step deformable b-spline syn with a 26-step b-spline distance). Linear and nonlinear warping matrices for fMRI-to-anatomical alignment were applied to individual scans in the time series, then the merged 4-D functional timeseries were moved to the atlas space using the prior anatomical-to-template transformation matrices.

A total of 64 ROI masks, divided into 32 left and 32 right ROI’s, were included in our analyses. Center voxel coordinates (see above) were used for 3D network visualizations in BrainNet viewer in MATLAB [24]. Signal timeseries were extracted from preprocessed fMRI scans with the assistance of ROI mask overlays. This generated 64 individual ROI text files per subject that contained L2-normalized resting state signals as a vector of 600 data points. The timeseries files were used in cross-correlations and in calculations of Pearson r coefficients for every pairwise combinations of ROIs (1dCorrelate in AFNI). The resulting number of pairwise correlations was 1,952 per subject (after removing 64 self-correlations). Correlation coefficients were imported to MATLAB and Fisher’s transform applied to ensure a normal distribution of z values prior to analyses.

### 2.5 Functional Network Analysis

Functional network analysis was completed as previously reported [25]. Briefly, the Brain Connectivity Toolbox [26] and MATLAB were used to determine weighted matrices. Edge densities thresholds were set in a range from 2 to 40% to calculate the following global network metrics: Clustering Coefficient (tendency of nodes to cluster and connect within the network), Characteristic Path Length (average of shortest path length between all pairs of nodes in the network), Transitivity (probability of neighboring nodes to be interconnected within the network), Global Efficiency (efficiency to communicate across distant brain regions), Louvain Modularity (density of connections within a neural cluster), and Small World Index (high local clustering with shortest path length). Unless otherwise indicated in the figures, a 10% threshold was used for all the node-specific measures such as Node Strength (how strongly a node directly connects to other nodes), Node degree (number of connections with other nodes), Eigenvector Centrality (node’s influence in a network), Participation Coefficient (node interaction within single or multiple communities), Local Efficiency (capacity to integrate information between neighboring nodes), and Betweenness Centrality (node’s influences over the flow of information in the network). Significant alterations in brain connectivity between sham and rmTBI were determined by using an unpaired T-test.

### 2.6 Brain volume and diffusion tensor imaging (DTI): Image Processing and Analysis

Anatomical T2-weighted images of sham and rmTBI mice were used to measure the brain volume using ITK-SNAP software. Diffusion weighted scans were acquired at 5-7 dpi using a 4-shot, 2-shell spin echo planar diffusion imaging (DTI EPI) sequence in Bruker Paravision, with TR = 4 seconds, TE = 19 ms, number of averages = 4, gradient duration δ= 3 ms, diffusion time Δ= 8 ms, 54 images with 3 different diffusion weightings, two b=0, 6 directions with b=600 s/mm^2^, and 46 directions with b=2000 s/mm^2^. A navigator signal was used by the Bruker reconstruction software to improve signal stability in the 4-shot EPI. Image saturation bands were placed on either side and below the brain to suppress non-brain signal during image acquisition. Diffusion images had the same FOV and slice thickness as the anatomical scan but with a lower resolution data matrix size of 128 × 96 and 17 slices (resolution: 0.117 mm x 0.117 mm x 0.7 mm) in the same space as anatomical scans. This allowed careful manual outlining of regions to be analyzed by using T2 scans and diffusion maps (see below).

Diffusion MRI scans were processed using tools available on FMRIB software library - FSL [27] and DSI Studio [28] as previously reported for mouse diffusion scans obtained at 11.1Tesla [29]. Individual diffusion images were examined carefully and removed from the image series due to low signal to noise or the presence of motion artifact. The gradient b-vector files were corrected by removing gradient directions for the removed images. Eddy correction was used for adjusting slight movements during image acquisition and gradient files rotated according to the motion correction vectors. After eddy correction, tensor element reconstruction and estimates of 1^st^, 2^nd^ and 3^rd^ eigenvectors and eigenvalues (λ1, λ2, λ3, where λ1 is axial diffusivity and the average of λ2 and λ3 provides radial diffusivity values) was performed using weighted least squares regression on DTIFIT in FSL [30]. This last step generated independent images of FA, mean, axial and radial diffusivities (MD, AD, and RD, respectively).

Regions of interest were manually selected in ITK-SNAP to estimate their mean intensity and volumes. These included thalamus, optic tract, fimbria, corpus callosum, hippocampus, and amygdala. Significant alterations in brain microstructure were determined by using an unpaired T-test.

### 2.7 Immunohistochemistry (IHC)

Formalin-fixed, Paraffin-embedded coronal brain sections (5 μm) were used for the IHC study. Both sham and rmTBI slides were processed simultaneously. The slides were deparaffinized in xylene twice for 5 min each and rehydrated sequentially in the gradient of ethanol followed by water for 3 min each. The antigen retrieval was performed by immersing slides in 10mM citrate with 0.5% tween20 (pH 6.0) and 30 min incubation in the steamer. The endogenous peroxidases were removed by incubating the slides with the mixture of 0.3%H2O2/PBS and 10% Triton X-100 (100μl for 200 ml of 0.3% H_2_O_2_ solution) for 20 min. Blocking was performed with 10% normal goat serum/PBS-T (0.05% tween) for 30 min at room temperature (RT). After blocking, the brain sections were incubated overnight with the following antibodies: Iba1 (1:1000, PA5-27436, Invitrogen) and GFAP (GA5) (1:1000, CS3670S, Cell Signaling) at 4°C for microglia and astrocytes, respectively. After incubation, the slides were washed with PBS-T and incubated for 30 min with corresponding biotinylated secondary antibody [Goat Anti-Rabbit IgG Antibody (H+L), Biotinylated-(BA-1000); Goat Anti-Mouse IgG Antibody, Biotinylated, R.T.U. BP-9200, Vector laboratories). Further, the sections were incubated with avidin-biotin complex (ABC) reagent for 20 min (VECTASTAIN Elite ABC-HRP Kit, Peroxidase Standard), PK-6100) following washes with PBS-T and developing with 3, 3’-diaminobenzidine (DAB) kit [KPL DAB Reagent Set, SeraCare (5510-0031)]. Then, the slides were washed twice in PBS, 5 min each, and counter-stained with hematoxylin for 1-2 min. The slides were washed briefly in water and dehydrated with the gradient of alcohol (70% ethanol, 90% ethanol and 100% ethanol) for 3 min each and incubated in xylene twice for 5 min each. The sections were mounted with Cytoseal Mountant 60 (Epredia 83104) and air dried. The slides were scanned using an Aperio image scanner (20X objective lens, Scan Scope™ XT, Aperio Technologies, Inc. Vista, CA, USA) and quantified using Image Scope software (v12.4.3.5008) positive pixel count program. Three sections per mouse at 55 μm apart were averaged to calculate the expression of Iba1 and GFAP.

### 2.8 Spatial proteomics analysis using the NanoString GeoMx Digital Spatial protein profiling platform

Formalin-fixed paraffin-embedded brain tissue was sectioned coronal at 5 μm thickness.

DSP staining and assay procedures were conducted according to the manufacturer-directed protocol. Briefly, to identify regions of interest (ROI), nuclei were fluorescently labeled with SYTO 13 (GeoMx Nuclear Stain Morphology kit, Item no. 121300303, Nanostring, Seattle, WA), and microglia cells were labeled with an Iba1 (GeoMx Alzheimer’s Morphology Kit, Item no. 121300306, NanoString, Seattle, WA, United States) antibody. The regions of interest (ROIs) were selected based on visual identification of anatomic regions using the nuclear stain, aided by increased Iba1 expression in white matter regions of injured animals. The following ROI were selected: primary somatic sensory area, dentate gyrus polymorph layer, thalamus, and the optic tract. The following protein cores and modules were used for multiplexed protein quantification: Neural Cell Profiling (Item no. 121300120), Parkinson’s disease (PD) pathology (Item no. 121300122), Alzheimer’s Disease (AD) pathology (Item no. 121300121), AD pathology extended (Item no. 121300123), Glial cell subtyping (Item no. 121300125), and autophagy (Item no. 121300124) markers. Samples were processed on the NanoString GeoMx system and the nCounter max system following manufacturer’s instructions. Data were analyzed using GeoMx DSP analysis suit Version 2.4.0.421 as suggested by manufacturer instructions. Briefly, the Field of view (FOV) QC was adjusted to 45% to check the low count score. All the QC passed segments were normalized to housekeepers such as GAPDH and Histone H3. Further background corrections were performed with the negative background targets such as Rb IgG, Rt IgG2a, and Rt IgG2b. background corrected files were changed to tab-delimited files, and custom scripts with slight modifications were used to generate volcano plots.

### 2.9 Statistical analyses

GraphPad Prism 9 software was used to perform all the statistical analysis. An unpaired T-test was used to determine significant differences in nodal properties between control and injured mice. Multiple t-tests were performed to measure the difference between density thresholds in rsfMRI data. An unpaired T-test was performed within the GeoMx DSP analysis suit to analyze the NanoString GeoMx Spatial Profiling data. All the results were represented as mean ± SD; p<0.05 is considered statistically significant.

## 3. Results

We used the Closed Head Injury Model of Engineered Rotational Acceleration (CHIMERA) that reproducibly causes DAI [11,31,32]. We injured 2.5-3mo C57Bl/6 mice, twice within a 24hr interval, and we measured outcomes after the primary injury phase (Fig. 1). As expected from clinical correlations with T2-weighted images, we found no overt structural or volumetric changes in the injured mice compared to controls (Fig. 2A-C).

**Fig. 1.**
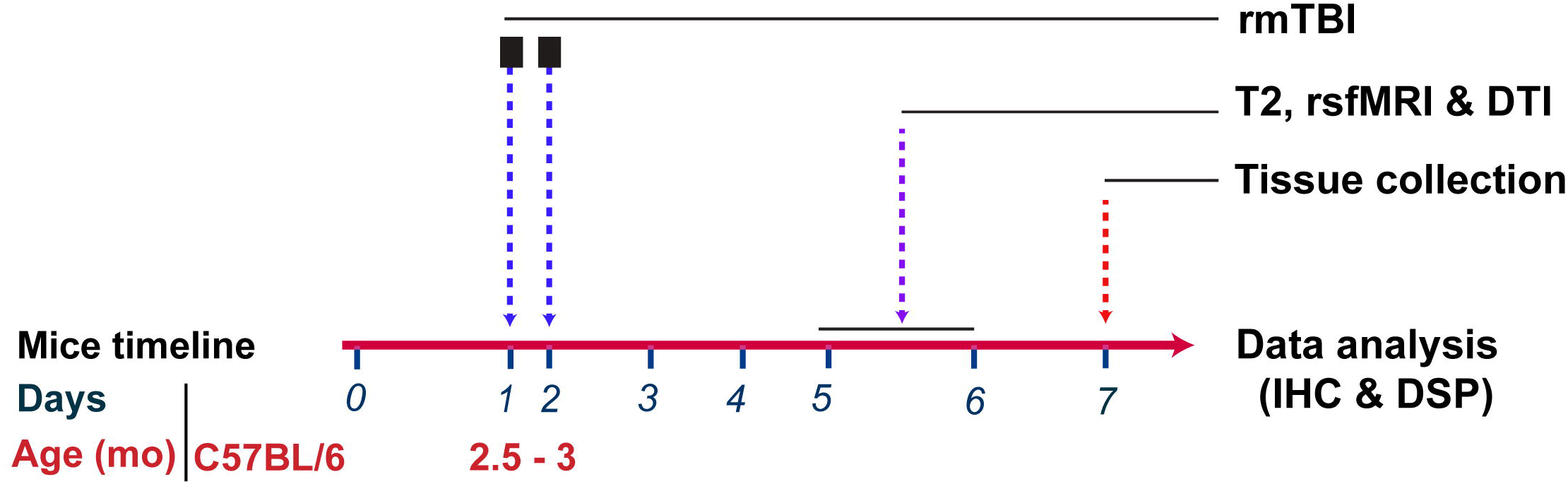
CHIMERA validation. The schematic representation and experimental design showing the number of rmTBIs, rsfMRI and tissue collection. Brains were collected at 7 dpi for Immunohistochemistry (IHC) and NanoString GeoMx spatial protein profiling analyses.

**Fig. 2.**
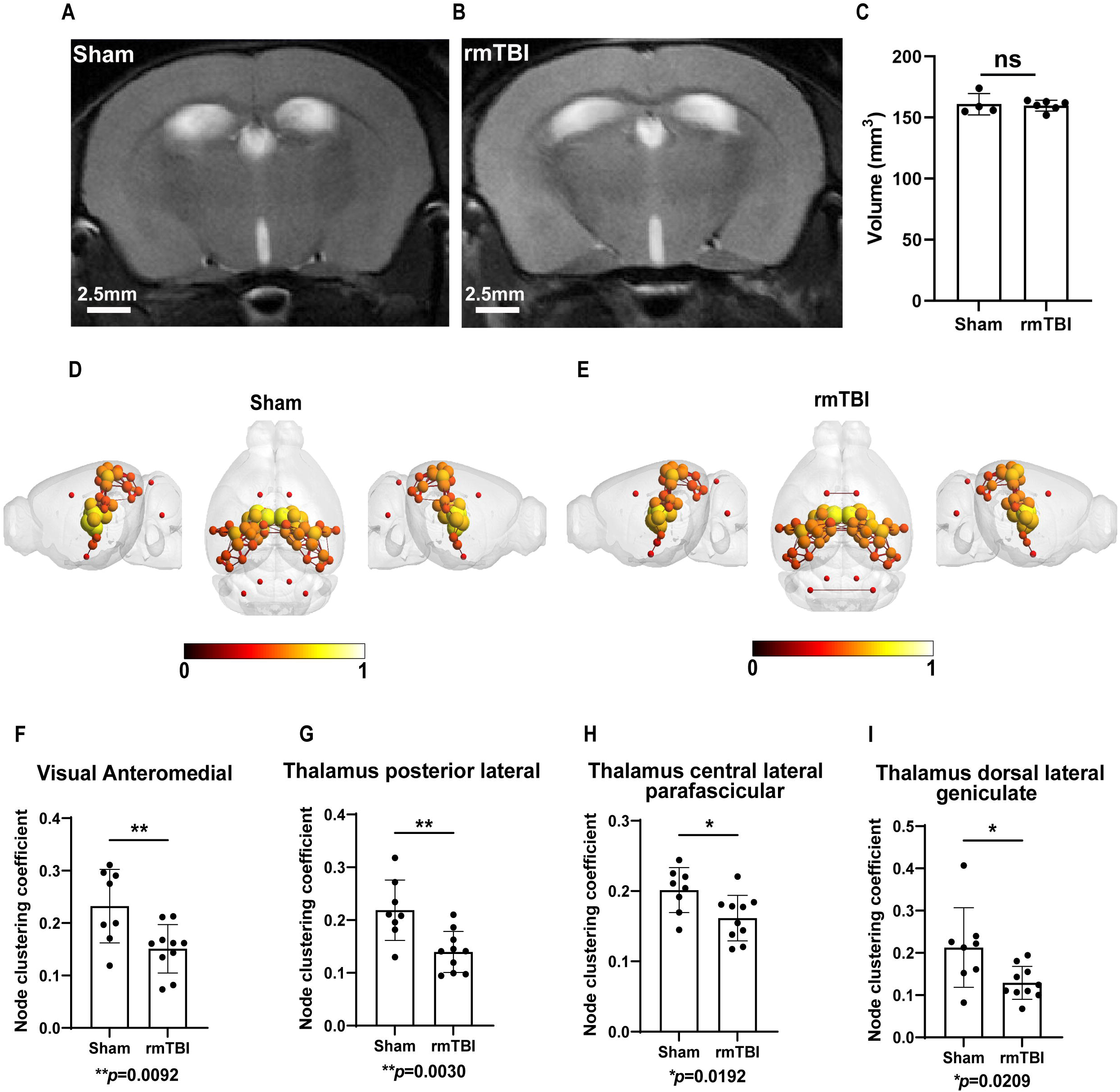
Clustering coefficient analysis reveals that the network integration and efficiency in specific brain regions are disrupted following rmTBI. Representative T2 weighted anatomical brain images of sham and rmTBI mice **(A&B)** were used for brain volume analysis. Quantification of brain volume revealed no changes between sham and rmTBI groups **(C)**. 3D functional connectome maps of sham and rmTBI mice brain shows the position of nodes and edges in the region of interest (ROIs) [total ROI = 64] **(D&E)**. Clustering coefficient in various regions of thalamus and visual area **(F-I)** were significantly altered in rmTBI group compared to sham. Unpaired t-test, mean ± SD, Sham (N= 8), rmTBI (N=10).

Given that changes in brain network organization strongly associate with cognitive dysfunction [33], we then coupled resting state functional MRI (rsfMRI) and graph theory network analyses to identify spatial and functional information following TBI [34]. We performed focal measurements in a rsfMRI modality such as node clustering coefficient, global and local efficiency, participation coefficient, eigenvector centrality, and betweenness centrality in specific brain regions that process visual, auditory, somatosensory information, fractional anisotropy (FA) and axial diffusivity in the optic tract; all in the absence of differences in brain volume. We developed graphs of functional brain networks by defining nodes (macroscopic regions of the brain) and edges (links). We selected 64 nodes (Supplementary table 1) that are closely associated with white matter tracts (Fig. 2D & E) for functional connectivity analysis. We calculated the node clustering coefficient, which is the index of the number of connected neighboring nodes [35]. We found that the clustering coefficient in specific thalamic and visual nodes were the most pronounced and significantly different changes identified (Fig. 2F-2I).

We then measured global efficiency and found that the edge density at 4% and 6% threshold was significantly decreased (*p*=0.024 and *p*=0.008, respectively) in the rmTBI group (Fig. 3A). Focusing on these densities, we found that local efficiency at the 4% (Fig. 3B-3C) and 6% (Fig. 3D-3F) density thresholds were significantly decreased in specific thalamic and visual nuclei. No changes were observed in the net clustering coefficient and small world index between sham and rmTBI groups (Fig. 3G-3H). In contrast, the net characteristic pathlength, revealing about functional network integration, was significantly increased (*p*=0.0367) in the rmTBI group at 4% edge density (Fig. 3I), but no changes were observed at the edge density ranges between 6-40%.

**Fig. 3.**
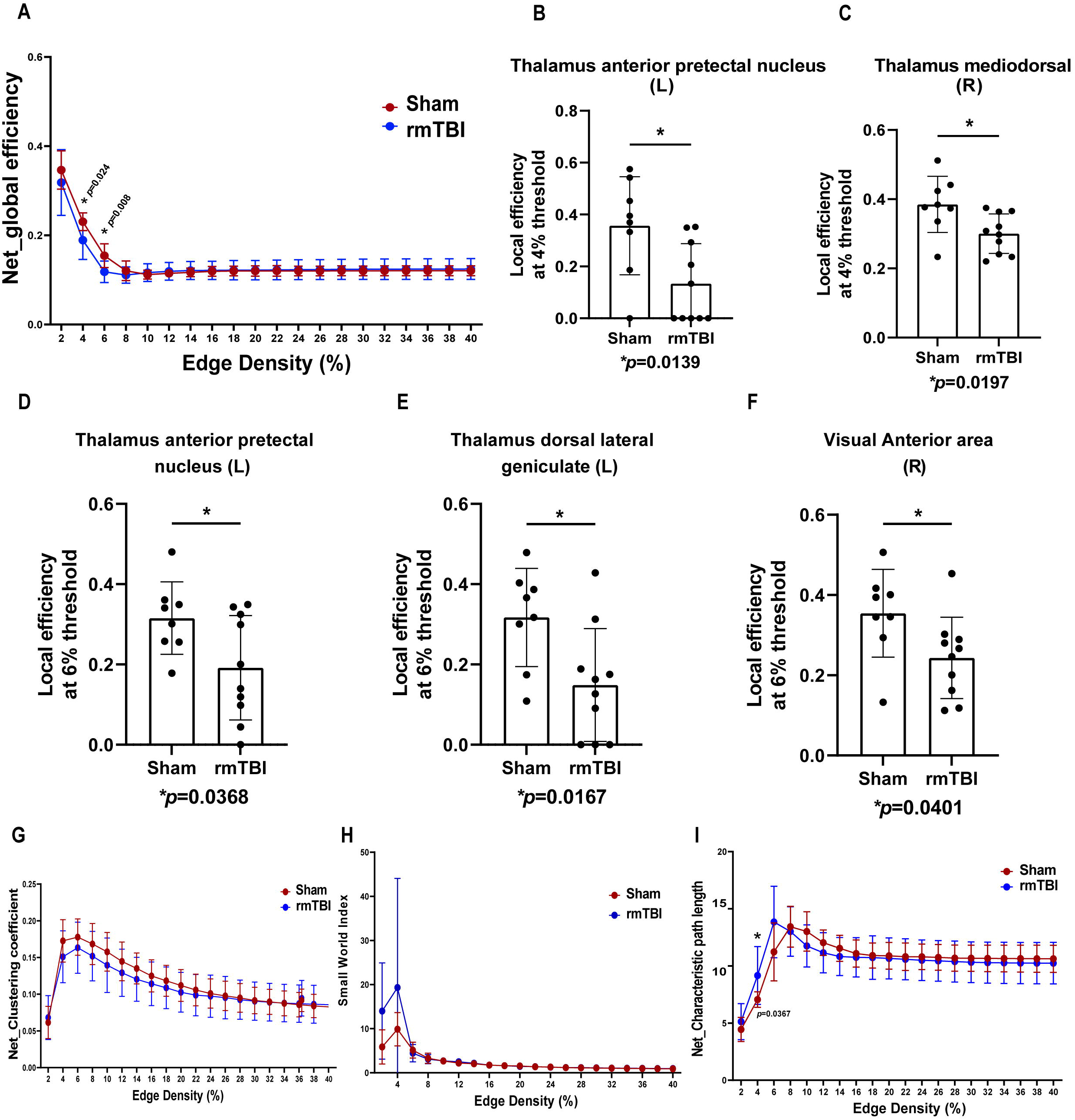
Global and Local efficiency analysis reveals that efficiency in specific brain regions are disrupted following rmTBI. Global efficiency **(A)** was significantly decreased at the density threshold of 4% and 6% in rmTBI group compared to Sham. Local efficiency at 4% threshold **(B & C)** was significantly decreased in thalamus anterior pretectal nucleus (L-Left) and thalamus mediodorsal (R-Right) regions of Injured mice. Similarly, Local efficiency at 6% threshold **(D-F)** was significantly decreased in thalamus anterior pretectal nucleus (L), thalamus dorsal lateral geniculate (L) and visual anterior area (R) of injured mice compared to Sham. No changes in global clustering coefficient **(G)** and small world index **(H)** were observed. The global characteristic pathlength **(I)** at 4% threshold was significantly increased in injured mice group compared to sham. The global metrices were analyzed by Multiple Unpaired t-test, mean ± standard error at edge densities ranges 2 to 40%. The groups comparison was analyzed by Unpaired t-test, mean ± SD, Sham (N= 8), rmTBI (N=10).

The greatest differences following rmTBI were detected in the strength of a node’s connections within its module (participation coefficient) and the influence of distinct nodes on the network (eigenvector centrality). We found that the participation coefficient in the left ventral posteromedial thalamic nuclei was significantly decreased (*p*=0.0026) in the rmTBI group (Fig. 4A). Conversely, participation coefficient was significantly increased in the right splenium of corpus callosum (*p*=0.0028), right central lateral parafascicular thalamic region (*p*=0.0137), right visual anteromedial area (*p*=0.0373), and right visual postrhinal area (*p*=0.0375) of the rmTBI group (Fig. 4B-4E). Eigenvector centrality in specific brain regions such as genu of corpus callosum (*p*=0.0166), the body (*p*=0.0329) and splenium of corpus callosum (*p*=0.0134), and the thalamic paracentral nucleus (*p*=0.0392) were significantly increased in the rmTBI group compared to sham (Fig. 4F-4I). The node betweenness centrality in the thalamic dorsal lateral geniculate nucleus was significantly increased in the rmTBI group (Fig. 4J). These highly unique characteristics reveal alterations to unique brain regions that define rmTBI node specificity. Meanwhile, other network features such as global network strength, Louvain modularity, assortativity, and Gamma values showed modest or undetectable differences between the sham and rmTBI groups. Only, the net lambda values, which indicate the degree of network randomness within neighboring nodes, was significantly decreased at wide edge densities from 8% to 40% (p<0.05) in rmTBI group (Fig. S1).

**Fig. 4.**
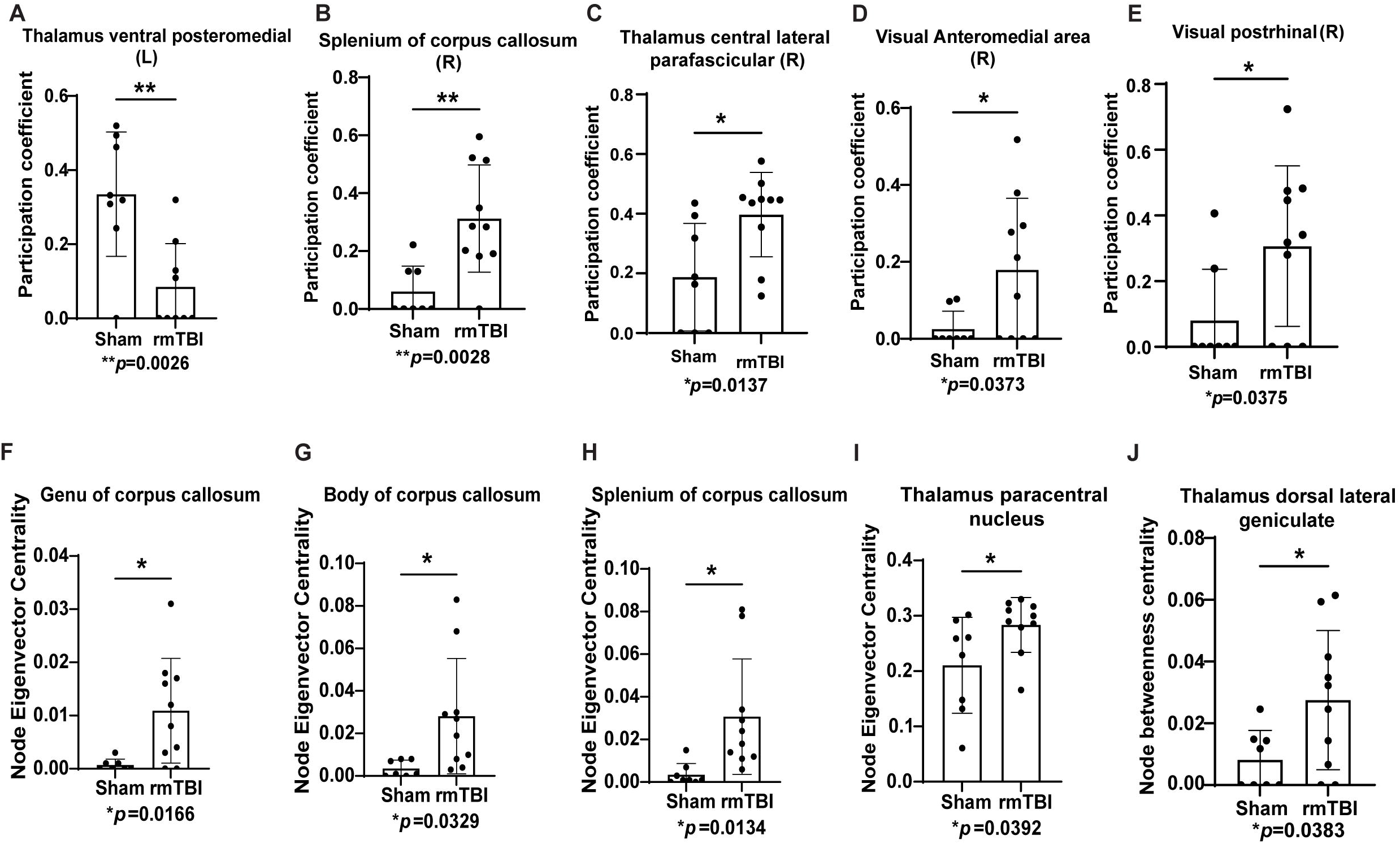
rmTBI disrupted connectivity between modules, increased eigenvector centrality and node betweenness centrality. Nodes in left (L) thalamus ventral posteromedial **(A)** region had low participation coefficient, but it is significantly increased in the right (R) splenium of corpus callosum, thalamus (R) and visual regions (R) of rmTBI group relative to control **(B-E)**. Increased eigenvector centrality shows increased high influence of specific nodes on connected networks following rmTBI in thalamic regions and corpus callosum **(F-I)**. The node betweenness centrality shows the nodes at the intersection between shortest path in the specific thalamus region **(J)** was significantly altered. Density threshold at which the participation coefficient and node eigenvector centrality were calculated. Unpaired t-test, mean ± SD.

Given FC differences in specific nodes, we examined cerebral microstructure by measuring the diffusion of water molecules in the thalamus, optic tract, and corpus callosum. We also focused on the fimbria, hippocampus, and amygdala because of their important role in learning and memory (Fig. 5A). We observed that the fractional anisotropy (FA), which measures white matter integrity, was significantly decreased (*p*=0.0240) in the optic tract of the rmTBI mice (Fig. 5B). The mean diffusivity (MD) (*p*=0.0259) in the hippocampus was significantly increased after rmTBI. A higher diffusion rate (increased water content) suggests the presence of edema or inflammation in this region (Fig. 5C). Two other measures of axonal integrity, radial diffusivity (RD; *p*=0.0303) and axial diffusivity (L1; *p*=0.0413) were also increased in the hippocampus of the rmTBI group, suggesting greater diffusion rate running parallel and perpendicular to white matter fiber tracts (Fig. 5D & 5E). As expected, due to the limited white matter content, we did not detect volumetric changes in the thalamus, fimbria, and amygdala between the sham and rmTBI groups.

**Fig. 5.**
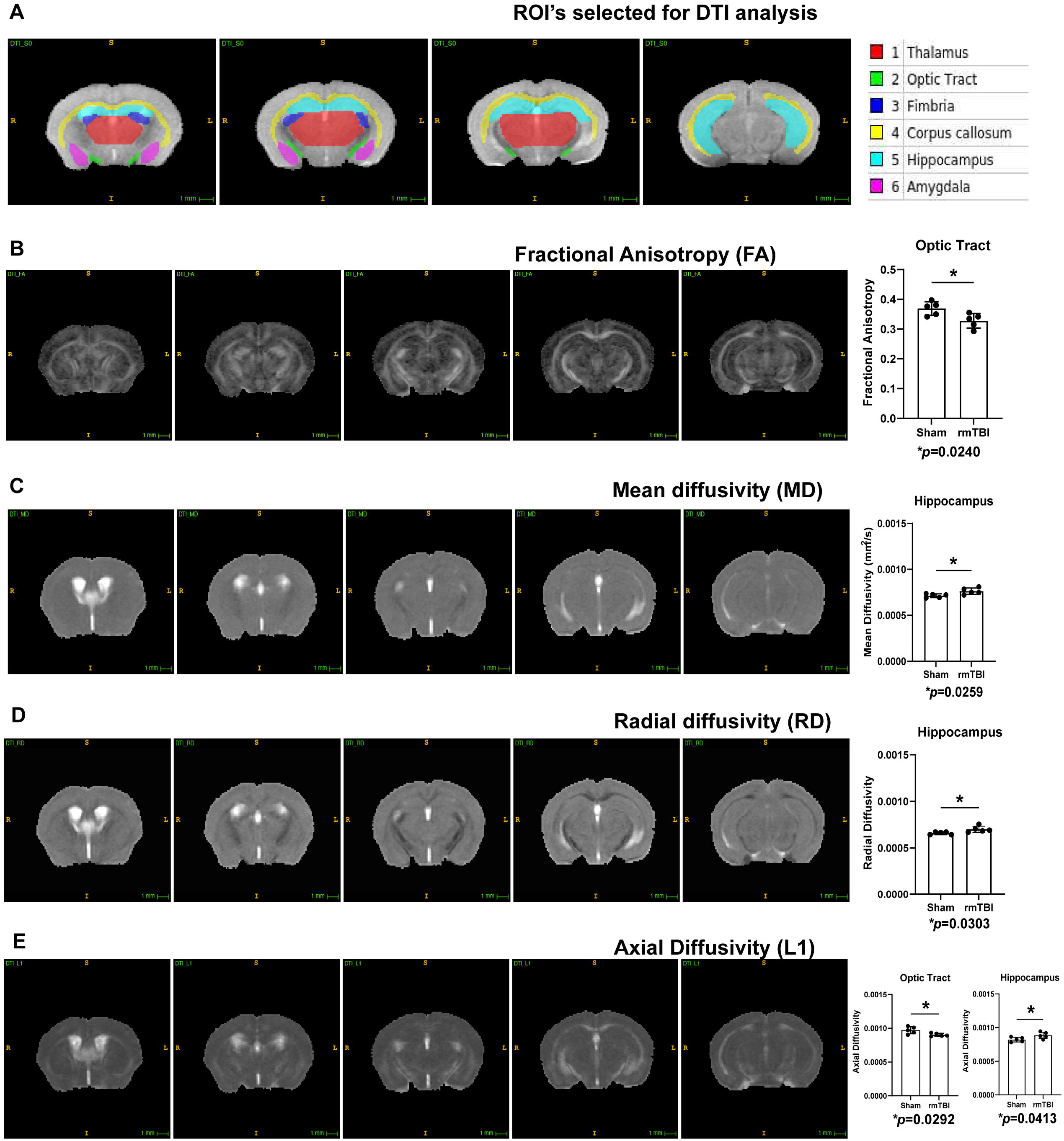
rmTBI altered tissue microstructure in specific brain regions. Representative DTI brain image shows **(A)** ROIs selected for microstructure pattern analysis. rmTBI increased fractional anisotropy (FA) in the optic tract **(B)**. Mean diffusivity (MD) and radial diffusivity (RD) were significantly increased in hippocampus of the injured mice **(C&D)**. Axial diffusivity (L1) was significantly increased in the optic tract and hippocampus of the injured mice **(E)** compared to sham. Unpaired t-test, mean ± SD.

We measured white matter gliosis by measuring localized changes in the levels of Iba1 and GFAP using immunohistochemistry. We detected a robust glial response in the optic tract (Fig. 6A-B and 6D-E) and corpus callosum (Fig. 6G-H) of injured mice. As established in previous reports, Iba1-positive microglia were significantly increased (*p*<0.0001) in the optic tract of injured mice (Fig. 6C). Similarly, GFAP-positive signal was significantly increased in both the optic tract (*p*<0.0001; Fig 6F) and corpus callosum (*p*<0.0320; Fig. 6I) of the rmTBI.

**Fig. 6.**
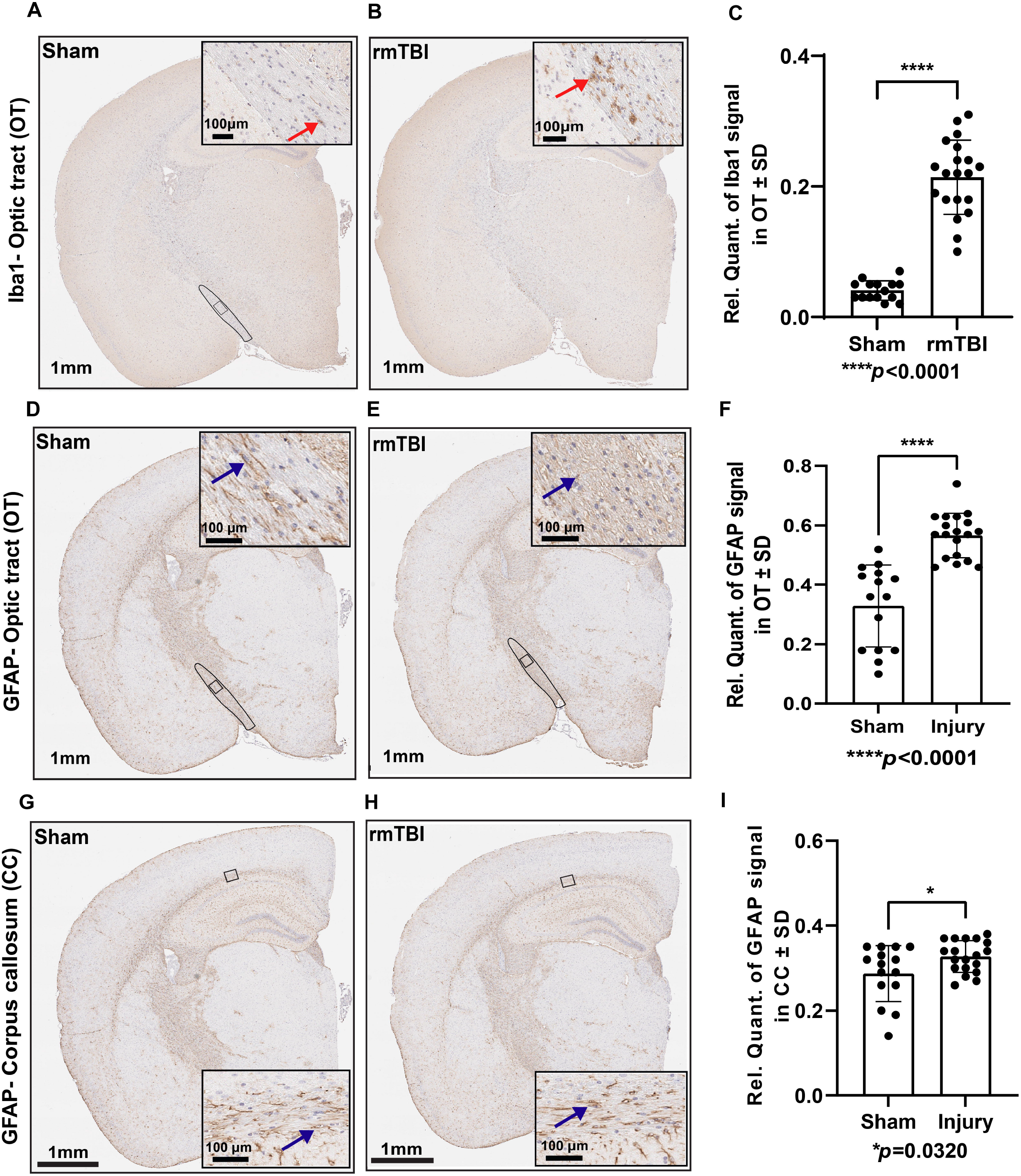
rmTBI induced gliosis in white matter tracts. Iba1 expression was significantly increased in the optic tract of injured mice compared to sham **(A-C)**. The red arrowhead indicates microglia. Brain coronal view: 1 mm; Scale bar for inset (enlarged view of the optic tract) = 100 μm. \*\*\*\**p*<0.0001, Unpaired t-test; Sham (N= 15), rmTBI (N=20). GFAP expression was significantly increased in the optic tract **(D-F)** and corpus callosum **(G-I)** of injured mice compared to sham. The blue arrowhead indicates astrocytes. Brain coronal view: 1 mm; Scale bar for inset (enlarged view of the optic tract and corpus callosum) = 100 μm. (\*\*\*\**p*<0.0001; \**p*= 0.0320, Unpaired t-test), Sham (N= 15), rmTBI (N=19).

The rsfMRI and DTI data identified four brain regions that were significantly altered by the injuries: the primary somatosensory area, hippocampus, thalamus, and optic tract. To determine whether the injuries impacted the levels of specific proteins in these areas, we performed digital spatial protein profiling using the NanoString GeoMx DSP platform. We measured changes in the levels of 72 proteins altered in ADRDs int the following ROI’s primary somatosensory area, dentate gyrus polymorph area, thalamus, and optic tract.

Levels of distinct proteins were significantly increased or decreased in the three areas of interest (Fig. 7A-B and Table 1). We observed no changes in the dentate gyrus polymorph layer (Data not shown). Myelin basic protein was significantly downregulated in the primary somatosensory area, which reflects white matter damage (Fig. 7N&O). Interestingly, both total tau and phospho-tau S396 (pS396) levels were significantly decreased but levels of one phospho-tau species, pS199 tau, were increased in the rmTBI group (Fig. 7C-7F). Various neuroinflammatory markers such as Iba1, GFAP, Cathepsin-D (Fig. 7H-7J) were significantly increased in the optic tract of the rmTBI group. The expression of neuroinflammatory marker GPNMB was also significantly increased, and cell proliferation marker Ki-67 was significantly decreased in the thalamic region (Fig. 7K-7M).

**Fig. 7.**
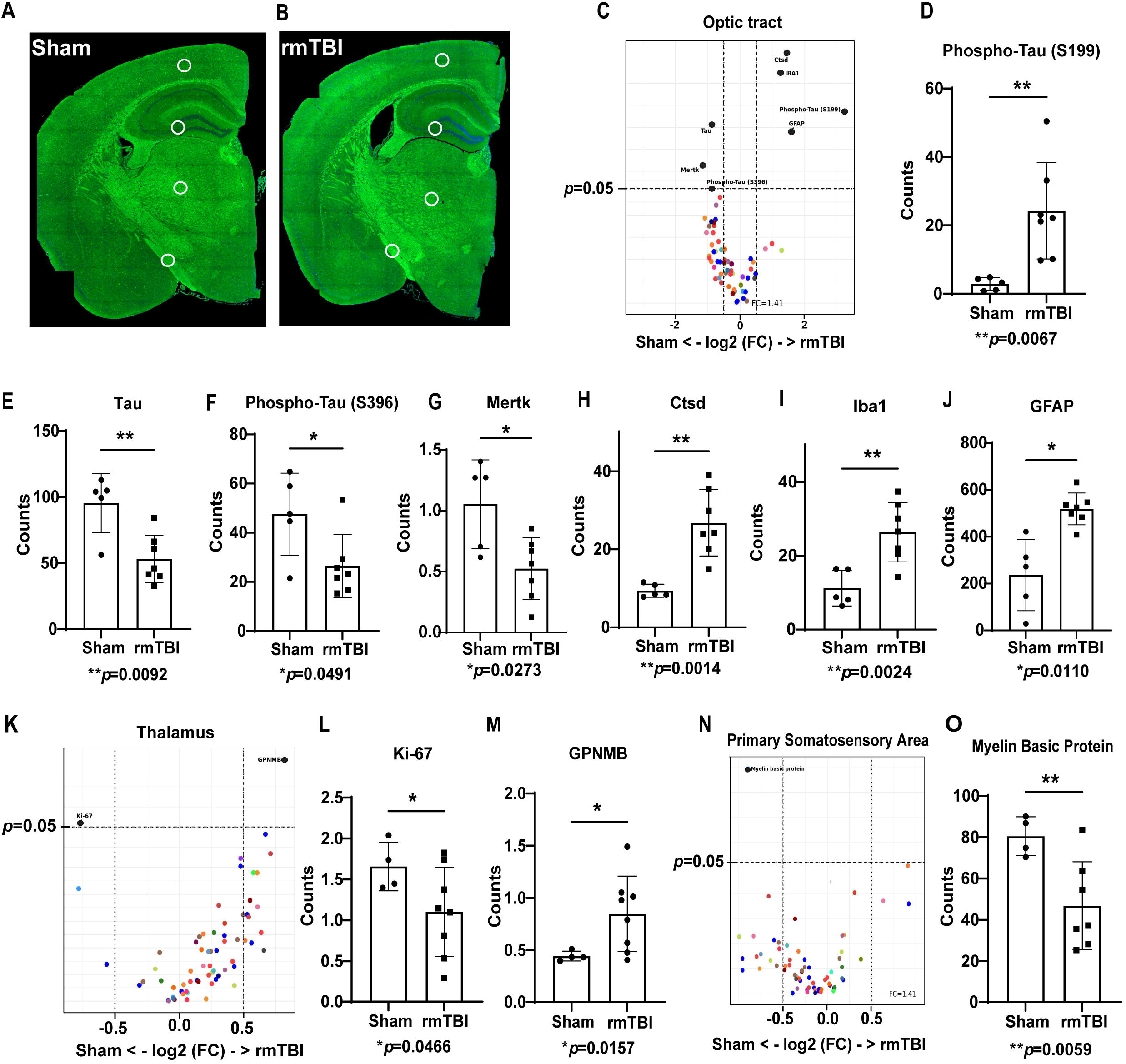
rmTBI increased the expression of disease associated proteins in white matter and grey matter regions. Coronal sections of sham and rmTBI mice shows the region of interest (primary somatic sensory area, dentate gyrus polymorph layer, thalamus, and the optic tract) selected for NanoString GeoMx spatial protein profiling analysis **(A&B)**. Volcano plots **(C)** and bar graphs **(D-J)** show significant alteration in the expression of Alzheimer’s disease (AD) pathological markers, glial and neural cell markers in the optic tract of the injured mice. Volcano plot **(K)** and bar graphs show **(L&M)** significant changes in the expression of cell proliferation and AD associated, neuroinflammatory markers in the thalamus of the injured mice. Myelin basic protein was significant decreased in the primary somatosensory area following rmTBI **(N&O)**. Protein levels were normalized to the expression of housekeeping proteins. Unpaired t-test, mean ± SD, N=4-8.

**Table. 1.**
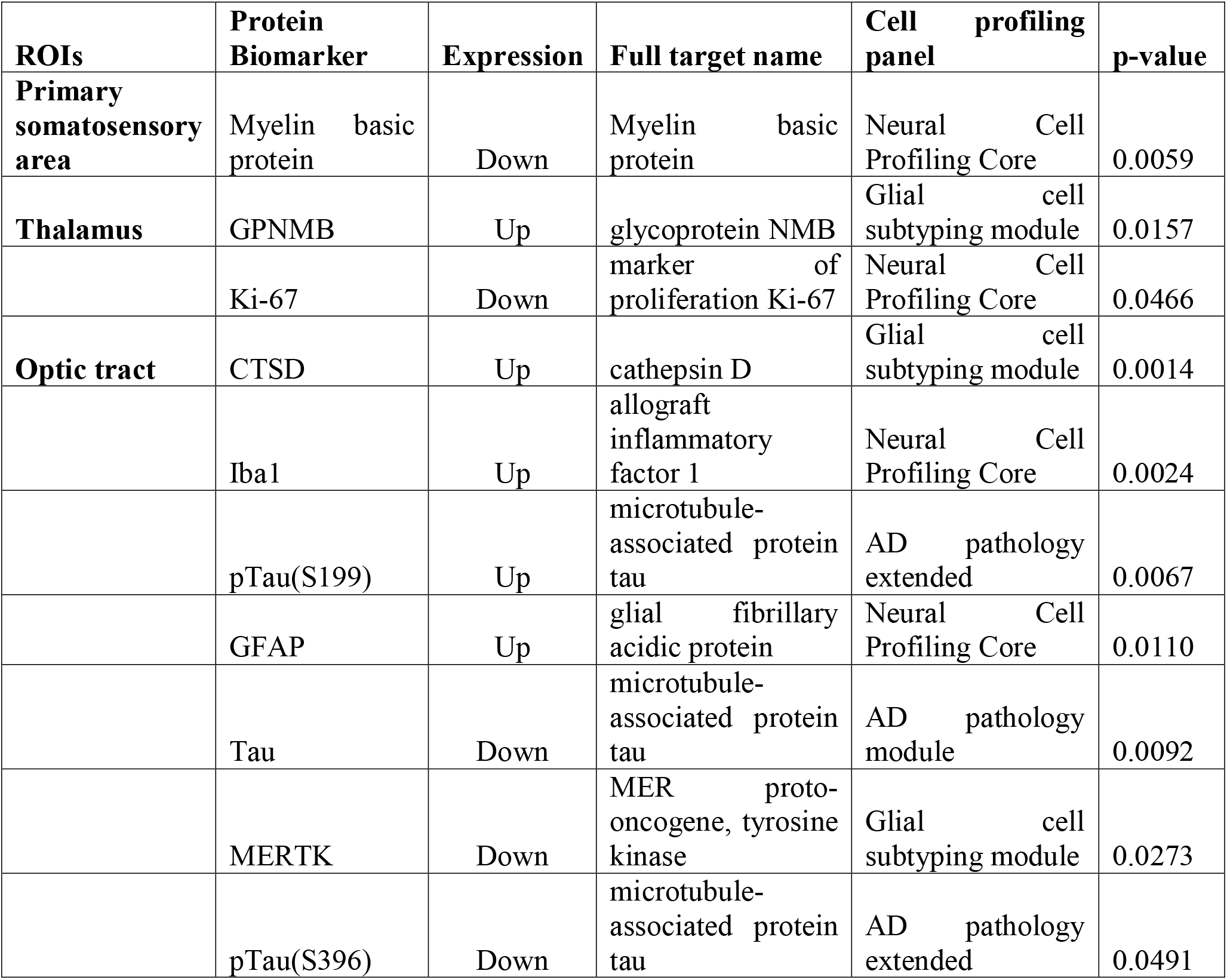
Summary of the expression of various neuropathological markers following rmTBI identified through NanoString GeoMx spatial protein profiling analysis.

These data align with both expected and previously reported alterations to molecular processes following head injury, including neuroinflammation (e.g. Iba1), white matter damage (e.g. MBP), and microtubule disruption (e.g. tau). Unexpectedly, changes in the levels of these proteins were unique to specific brain regions, suggesting that the injury-induced alterations unleash distinct consequences. Importantly, these distinct changes could be the root of the imaging alterations identified in the rsfMRI, and they may serve as biomarkers that distinguish clinical outcomes.

## 4. Discussion

The long-term goal of this study was to establish novel mechanistic information that connects head injuries to neurodegenerative disorders. Here, we identified rmTBI-induced imaging abnormalities and aberrant accumulation of unique proteins. We determined that injuries impact brain regions in distinct manners, and these distinctions may explain why rmTBI manifests differently between subjects. We also identified proteins that are increased or decreased depending on the brain region (Table 1). Our data strongly suggest that the molecular pathways leading to accumulation or depletion of these proteins contributes to the imaging abnormalities we identified.

Investigating brain networks using rsfMRI and graph theory offers powerful functional measures, all of which can be translated to the clinic. Network alterations, such as the ones described herein, have strong diagnostic potential for TBI among other brain disorders [36–38]. Recent work using rsfMRI identified broad functional connectivity deficits in human subjects; these defects correlated with diminished choice-reaction scores, showing high sensitivity. Our work improves upon pre-clinical and clinical studies because we dissected more granular aspects of functional connectivity in a mouse model of rmTBI and at ultra-high field [25,39–41]. Functional graph metrics such as clustering coefficient, characteristic path length, centrality, and efficiency are quantifiable features that reveal how information flows throughout the network [42], and they were impaired following rmTBI (Fig. 2–4). Our findings are concordant with other pre-clinical models of mild head injury showing local alterations in betweenness centrality, clustering coefficient, and local efficiency, while global changes were intact [43], showing that these robust measures could be applicable to milder and [44,45] repetitive TBI.

Although we did not detect global changes (as reported in Boroda et al., 2021[46]) in clustering coefficient, the local clustering coefficient at 10% edge density was significantly reduced in the visual anteromedial area and various sub-regions of the thalamus in the rmTBI group. Further, we observed a significant decrease in net global and local efficiency in the rmTBI group at thresholds below 10% densities which contain true connections; this measure is commonly used in clinical practice [44,45]. These results positively reveal that local integration, efficiency, and integrative information processing across distal brain regions were significantly altered following rmTBI. Consistent with previous reports, we did not observe changes in the small-world index and net clustering coefficient after mTBI [25].

We also calculated eigenvector and betweenness centrality to assess the highly influential nodes and nodes at the intersection between the shortest path, respectively [25]. We observed high betweenness centrality scores in thalamus dorsal lateral geniculate and high eigenvector centrality scores in modules with nodes located in sub-regions of corpus callosum and thalamus. Our results indicate that the injury increased the distance between connections of the thalamus and corpus callosum thereby impacting the connectivity between clusters in these networks. We note that these nodal changes are not observed in all TBI models. Yang et al., 2021[25] reported no differences in betweenness centrality in Controlled cortical impact (CCI) rats at 30 dpi. Given that nodes in eigenvector centrality scores in the contralateral cortex can be restored from 2 to 30 days after injury, it will be important to assess these metrics at later post-injury periods to confirm the long-term repercussions of rmTBI. Confirming these changes at different timepoints will provide valuable information about compensatory mechanisms and reorganization of damaged networks after TBI.

The thalamus is an important deep gray matter region of the brain composed a bundle of white matter tracts and groups of nuclei, which plays a vital role in global multifunctional pathways. It is highly susceptible to damage due to sudden acceleration or deceleration movements during head trauma [38]. Consistent with previous reports, our results show that rmTBI induced by CHIMERA damages thalamic nuclei. Given its numerous roles in integrating brain signaling, damage to the thalamus may result in varied manifestations after TBI; this may underlie why patients present broad symptomatology.

In addition to the network-associated changes, aberrant signaling at the molecular level is a probable pathogenic factor in the transition from rmTBI to chronic neurodegeneration. We observed significant changes in distinct proteins that aligned with unique brain regions. For example, significant accumulation of Iba1 and GFAP in the optic tract, which was consistent with the immunohistochemistry results and previous studies using CHIMERA [11], indicate that rmTBI induced gliosis. In addition, glycoprotein non-metastatic melanoma protein B (GPNMB) was significantly increased in the thalamus of the rmTBI group. GPNMB is highly expressed in glia, and is linked with risk for Parkinson’s disease [47,48].

Given their limited and unclear molecular links with TBI, follow-up experiments evaluating the prognostic value of GPNMB, Ki67, and cathepsin-D after rmTBI could unveil new biomarkers. Interestingly, total and pS396 tau levels were decreased in the optic tract were decreased, while pS199 tau was increased. We hypothesize that this could be the result of tau cleavage resulting from head injury; importantly, it suggests that distinct post-translationally modified tau species could extravasate and reflect the state of injuries thereby serving as more reliable biomarkers [7]

The outcomes of the present study provide important information about the effect of rmTBI and its primary molecular mechanism on the underlying pathology at 5-7dpi; this period corresponds to secondary injury, where complex molecular pathways are unleashed after injury. Importantly, changes in this time point could precede the long-term consequences of rmTBI and connect injuries with the pathophysiology of TBI-associated neurodegenerative diseases. Our future work on the assessment of the post-TBI effect together with behavioral outcomes will help to understand the molecular mechanisms of TBI and their long-term consequences on cognition.

## 5. Conclusions

Using highly translational imaging modalities and digital spatial protein profiling we report abnormalities following CHIMERA-induced rmTBI 5-7dpi. These measures may serve as potential biomarkers that can diagnose rmTBI and predict clinical outcomes. Moreover, protein changes could be responsible for the molecular pathways involved in the acute and chronic consequences of rmTBI. Studies to identify the longevity of these abnormalities and their association with cognitive dysfunction are currently underway.

## Acknowledgements

This work was primarily supported by grant 1 R01 AG074584-01 from NIH/NIA. A portion of this work was performed in the McKnight Brain Institute at the National High Magnetic Field Laboratory’s Advanced Magnetic Resonance Imaging and Spectroscopy (AMRIS) Facility, which is supported by National Science Foundation Cooperative Agreement DMR-1644779 and the State of Florida. This work was supported in part by an NIH award, S10 RR025671, for MRI/S instrumentation.

## Supplementary information Legend

**Fig. S1. Network disruption by rmTBI is not widespread**. No differences in the Network strength **(A)**, Louvain modularity **(B)**, Assortativity **(C)**, Transitivity **(D)** and Gamma **(E)**. Net lambda was significantly decreased in rmTBI group at various density threshold levels compared to sham **(F)**. Multiple unpaired t-test, mean error ± SD at edge densities ranges 2 to 40%.

## Supplementary table Legend

**Supplementary table. 1.** The complete list of 64 ROIs (left and right) analyzed through rsfMRI and graph theory.

| **ROI #** | **Abbreviation** | **Region** |
| --- | --- | --- |
| 1 | L VISAA | Visual Anterior area |
| 2 | L VISRLA | Visual Rostrolateral area |
| 3 | L VISAMA | Visual Anteromedial area |
| 4 | L VISLA | Visual Lateral area |
| 5 | L VISPV1 | Visual primary visual |
| 6 | L VISPV2 | Visual primary visual |
| 7 | L VISPL | Visual Posterolateral |
| 8 | L VISPM | Visual Posteromedial |
| 9 | L VISPR | Visual postrhinal |
| 10 | L THA MED | Thalamus mediodorsal |
| 11 | L THA L | Thalamus anterolateral complex of ventral thalamus |
| 12 | L THA AC | Thalamus laterodorsal |
| 13 | L THA PC | Thalamus posterior complex |
| 14 | L THA VP | Thalamus ventral posteromedial |
| 15 | L THA para | Thalamus paracentral nucleus |
| 16 | L THA PL | Thalamus posterior lateral |
| 17 | L THA CL | Thalamus central lateral – parafascicular |
| 18 | L THA AP | Thalamus anterior pretectal nucleus |
| 19 | L THA DL | Thalamus dorsal lateral geniculate |
| 20 | L THA V | Thalamus ventromedial |
| 21 | L TEMP | Temporal association areas |
| 22 | L DEN | dentate nuclei of cerebellum |
| 23 | L CERE | interposed complex of cerebellum |
| 24 | L CP G | genu of corpus callosum |
| 25 | L CP1 | Body of corpus callosum 1 |
| 26 | L CP2 | Body of corpus callosum 2 |
| 27 | L SCP1 | splenium of corpus callosum 1 |
| 28 | L SCP2 | splenium of corpus callosum 2 |
| 29 | L SCP3 | splenium of corpus callosum 3 |
| 30 | L OT1 | Optic tract 1 |
| 31 | L OT2 | Optic tract 2 |
| 32 | L OT3 | Optic tract 3 |
| 33 | R VISAA | Visual Anterior area |
| 34 | R VISRLA | Visual Rostrolateral area |
| 35 | R VISAMA | Visual Anteromedial area |
| 36 | R VISLA | Visual Lateral area |
| 37 | R VISPV1 | Visual primary visual |
| 38 | R VISPV2 | Visual primary visual |
| 39 | R VISPL | Visual Posterolateral |
| 40 | R VISPM | Visual Posteromedial |
| 41 | R VISPR | Visual postrhinal |
| 42 | R THA MED | Thalamus mediodorsal |
| 43 | R THA L | Thalamus anterolateral complex of ventral thalamus |
| 44 | R THA AC | Thalamus laterodorsal |
| 45 | R THA PC | Thalamus posterior complex |
| 46 | R THA VP | Thalamus ventral posteromedial |
| 47 | R THA para | Thalamus paracentral nucleus |
| 48 | R THA PL | Thalamus posterior lateral |
| 49 | R THA CL | Thalamus central lateral – parafascicular |
| 50 | R THA AP | Thalamus anterior pretectal nucleus |
| 51 | R THA DL | Thalamus dorsal lateral geniculate |
| 52 | R THA V | Thalamus ventromedial |
| 53 | R TEMP | Temporal association areas |
| 54 | R DEN | dentate nuclei of cerebellum |
| 55 | R CERE | interposed complex of cerebellum |
| 56 | R CP G | genu of corpus callosum |
| 57 | R CP1 | Body of corpus callosum 1 |
| 58 | R CP2 | Body of corpus callosum 2 |
| 59 | R SCP1 | splenium of corpus callosum 1 |
| 60 | R SCP2 | splenium of corpus callosum 2 |
| 61 | R SCP3 | splenium of corpus callosum 3 |
| 62 | R OT1 | Optic tract 1 |
| 63 | R OT2 | Optic tract 2 |
| 64 | R OT3 | Optic tract 3 |

