## Supplementary figures and images for "Repetitive mild traumatic brain injury impairs resting state fMRI connectivity and alters protein profile signaling networks"

### Fig. S1

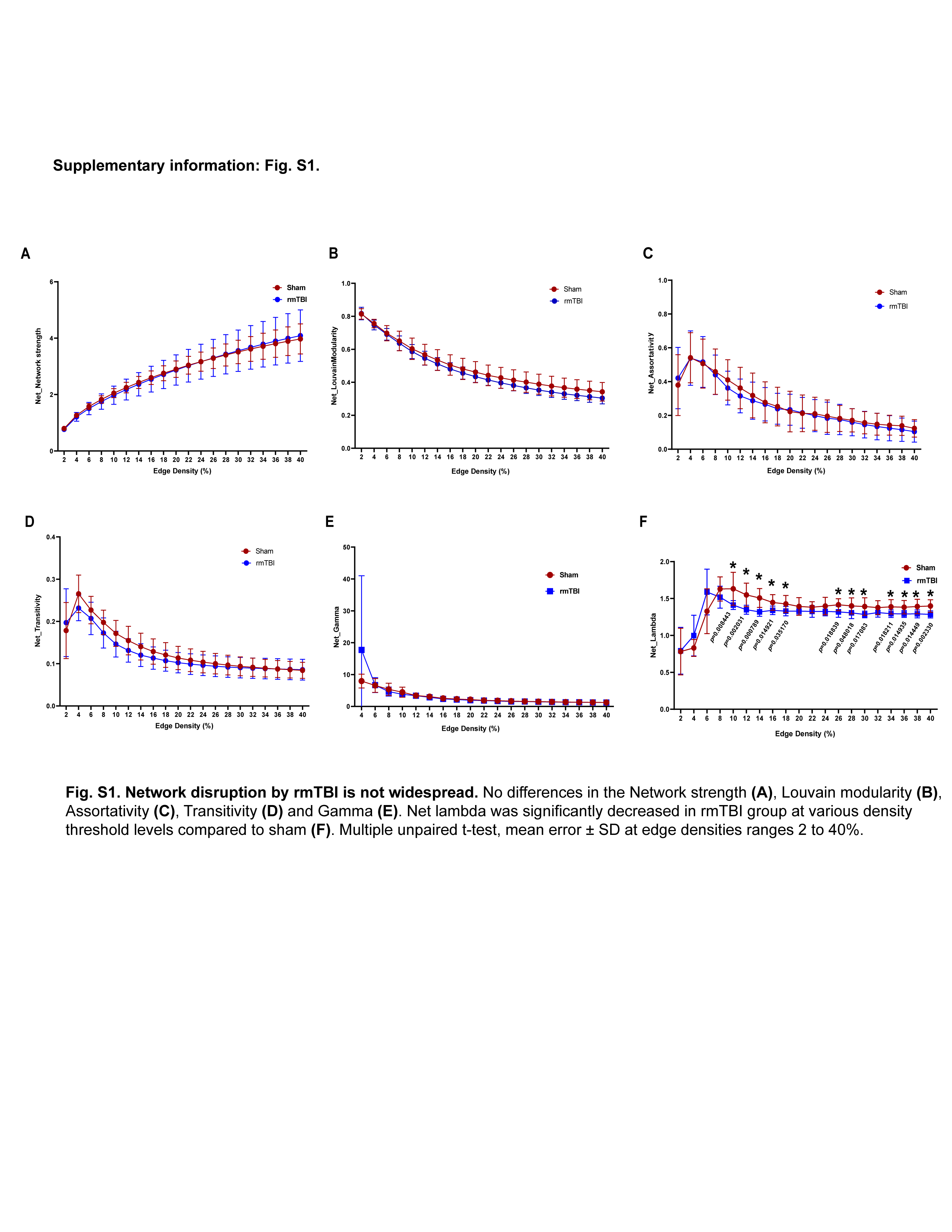
